# MetAP2 inhibition reduces food intake and body weight in a ciliopathy mouse model of obesity

**DOI:** 10.1101/800151

**Authors:** Tana S. Pottorf, Micaella Fagan, Bryan Burkey, David J. Cho, James E. Vath, Pamela V. Tran

## Abstract

The ciliopathies Bardet-Biedl Syndrome and Alström Syndrome are genetically inherited pleiotropic disorders with primary clinical features of hyperphagia and obesity. Methionine aminopeptidase 2 inhibitors (MetAP2i) have been shown in preclinical and clinical studies to reduce food intake, body weight, and adiposity. Here we investigated the effects of MetAP2i administration in a mouse model of ciliopathy produced by conditional deletion of the *Thm1* gene in adulthood (*Thm1* cko). *Thm1* cko mice show decreased hypothalamic *pro-opiomelanocortin* expression as well as hyperphagia, obesity, metabolic disease and hepatic steatosis. In obese *Thm1* cko mice, two-week administration of MetAP2i reduced daily food intake and reduced body weight 17.1% from baseline (vs. 5% reduction for vehicle). This was accompanied with decreased levels of blood glucose, insulin and leptin. Further, MetAP2i reduced gonadal adipose depots and adipocyte size and improved liver morphology. This is the first report of MetAP2i reducing hyperphagia and body weight, and ameliorating metabolic indices in a mouse model of ciliopathy. These results support further investigation of MetAP2 inhibition as a potential therapeutic strategy for ciliary-mediated forms of obesity.

## Introduction

Obesity and associated insulin resistance increase risk for potentially fatal chronic diseases, including cardiovascular disease, type 2 diabetes, and non-alcoholic fatty liver disease. As obesity is pandemic^1^, finding effective therapeutic strategies is critical to improving global health.

In preclinical and clinical studies, inhibition of methionine aminopeptidase 2 (MetAP2) has shown promising results. MetAP2 belongs to a family of metalloproteases, which cleaves the N-terminal methionine of nascent proteins. This post-translational modification induces subcellular localization changes and activation of the targeted protein^2^. Fumagillin, a natural product of *Aspergillus fumigatus*, irreversibly inhibits MetAP2^3^. MetAP2 inhibition causes late G1 cell cycle arrest, and inhibits cell proliferation, as well as phosphorylation of extracellular signal-regulated kinase 1 and 2 (ERK½)^4–7^. Early studies revealed anti-cancer and anti-fungal effects of fumagillin^8,9^. Subsequently, studies showed fumagillin and its analogs also have anti-obesity effects, resulting in decreased body weight and adiposity and increased insulin sensitivity in high-fat diet-induced obese mice and rats^10,11^, as well as in genetic *ob/ob* mice^12^. Additionally, administration of the fumagillin derivative, beloranib, to individuals with non-genetic causes of obesity and to patients with Prader-Willi Syndrome, a genetic disease that causes insatiable appetite and obesity, resulted in reduced food intake and body weight^13,14^. The effectiveness of MetAP2i in various models raises the possibility that inhibiting MetAP2 may counter other forms of obesity.

Ciliopathies are genetic disorders that arise from dysfunctional or absent cilia, and present numerous clinical features, including renal and hepatic fibrocystic disease, skeletal defects, infertility, hydrocephalus, mental disability, brain malformations, and central obesity^15^. Primary cilia are microtubule-based, mechanosensory organelles that protrude from the apical membrane of most mammalian cells and regulate signaling pathways. Primary cilia utilize intraflagellar transport (IFT) multi-protein complexes for bi-directional movement of protein cargo along the ciliary axoneme. The IFT-B complex mediates anterograde protein transport, while the IFT-A complex is required for retrograde transport and for ciliary import of membrane-associated and signaling proteins^16,17^. Another multi-protein complex, the BBSome, transports signaling molecules to the ciliary base, and acts like an adaptor between IFT complexes and protein cargo in the ciliary export of signaling molecules. Two ciliopathies, Alström Syndrome and Bardet-Biedl Syndrome (BBS), present obesity as a central clinical feature^18,19^. Additionally, polymorphisms in the *BBS* genes in the general population have been associated with obesity, and cilia length defects have been identified in adipose-derived mesenchymal stem cells from obese individuals, suggesting a more common relevance for cilia-related mechanisms^20–22^.

Modifying mutations in the IFT-A gene, *THM1*, have been reported in patients with Bardet Biedl Syndrome^23^. We have shown that global deletion of *Thm1* in adult mice causes decreased hypothalamic expression of appetite-controlling *pro-opiomelanocortin* (*Pomc*), hyperphagia, obesity and metabolic syndrome^24^. Here we examine the effects of administering a fumagillin derivative to obese *Thm1* conditional knock-out (cko) mice. Our results reveal reduced food intake, body weight and adipose tissue mass, as well as improved metabolic indices. These data indicate MetAP2 inhibition as a potential therapeutic strategy against obesity caused by genetic disorders of cilia.

## Results

### MetAP2i treatment decreases body weight and food intake in *Thm1* cko mice

To generate obese *Thm1* cko mice, we induced deletion of *Thm1* in male mice at five weeks of age and fed mutant mice and control littermates *ab libitum* throughout the 13-week duration of the study (Fig 1A). Body weight was measured weekly from 0-10 weeks post-*Thm1* deletion. At 10 weeks following gene deletion, *Thm1* cko mice weighed 42.2g ± 1.1g, while control littermates weighted 29.4g ± 0.7g, confirming the obese phenotype of the mutant mice (Fig S1A). At this timepoint (week 10 of the experiment), mice were housed individually and baseline measurements of body weight and food intake were obtained daily for one week. From week 10-11 of the experiment, *Thm1* cko mice showed a higher average daily food intake than control littermates, consistent with hyperphagia (Fig 1B). Unexpectedly, we observed that individual housing induced weight loss in many of the control mice and in all of the *Thm1* cko mice, and that the mutant animals showed a greater percent weight reduction (−3.8%) than control animals (−0.8%; Fig 1C; Fig S1B). Following this one-week observation period, we administered daily subcutaneous injections of MetAP2i or vehicle for two weeks (week 11-13). MetAP2i treatment reduced food intake relative to vehicle in *Thm1* cko mice (Fig 1D). Additionally, MetAP2i treatment caused a −17.1% body weight reduction compared to −5.0% for vehicle in *Thm1* cko mice (Fig 1D; Fig S1C). These data show that MetAP2i can counter the hyperphagia and increased body weight induced by deletion of *Thm1*.

**Figure 1.**
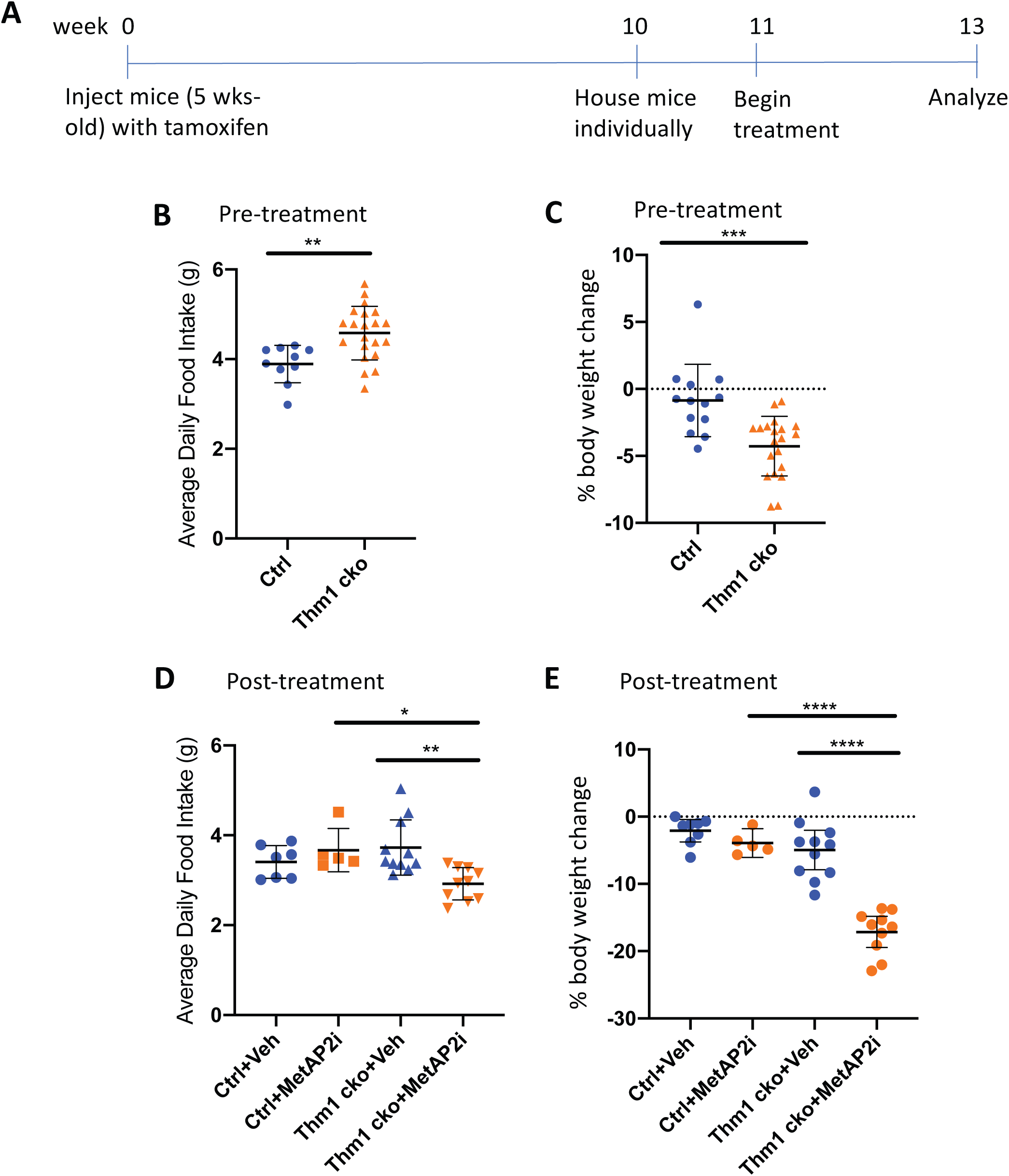
MetAP2i treatment decreased food intake and body weight in *Thm1* cko mice. **A)** Experimental timeline. Mice were injected at 5 weeks of age (week 0 of experiment) with tamoxifen to induce deletion of *Thm1*. Mice were fed *ab libitum* throughout the experiment. At week 10 (10 weeks post-tamoxifen injection), the obese phenotype was ascertained in *Thm1* cko mice, then mice were housed individually and food intake and body weight were measured daily until the end of the experiment. From week 11-13, daily subcutaneous injections of MetAP2i or vehicle were administered. **B)** Average daily food intake from week 10-11. **C)** Percent body weight change from week 10-11. **D)** Average daily food intake from week 11-13. **E)** Percent body weight change from week 11-13. *P<0.05; **P<0.005; ***P<0.0005; ****P<0.00005

### MetAP2i treatment improved metabolic indices in *Thm1* cko mice

Following the two-week intervention, we measured metabolic parameters, including non-fasting blood glucose and serum insulin and leptin, which we previously observed to be elevated in obese *Thm1* cko mice^24^. In this study, *Thm1* cko mice did not have higher non-fasting blood glucose than control littermates (Fig 2A). Yet in *Thm1* cko mice, MetAP2i treatment reduced non-fasting blood glucose levels relative to vehicle treatment (Fig 2A). Serum insulin levels were higher in vehicle-treated *Thm1* cko mice than in vehicle-treated control littermates, but MetAP2i treatment in *Thm1* cko mice decreased insulin levels relative to vehicle treatment and to an extent that insulin levels were not significantly different from vehicle-treated control mice (Fig 2B). Similarly, serum leptin was elevated in vehicle-treated *Thm1* cko mice relative to control littermates. In *Thm1* cko mice, MetAP2i treatment reduced leptin levels relative to vehicle treatment and to a degree that leptin levels were not significantly different from those of vehicle-treated control mice (Fig 2C). These data indicate that MetAP2i can correct these metabolic parameters.

**Figure 2.**
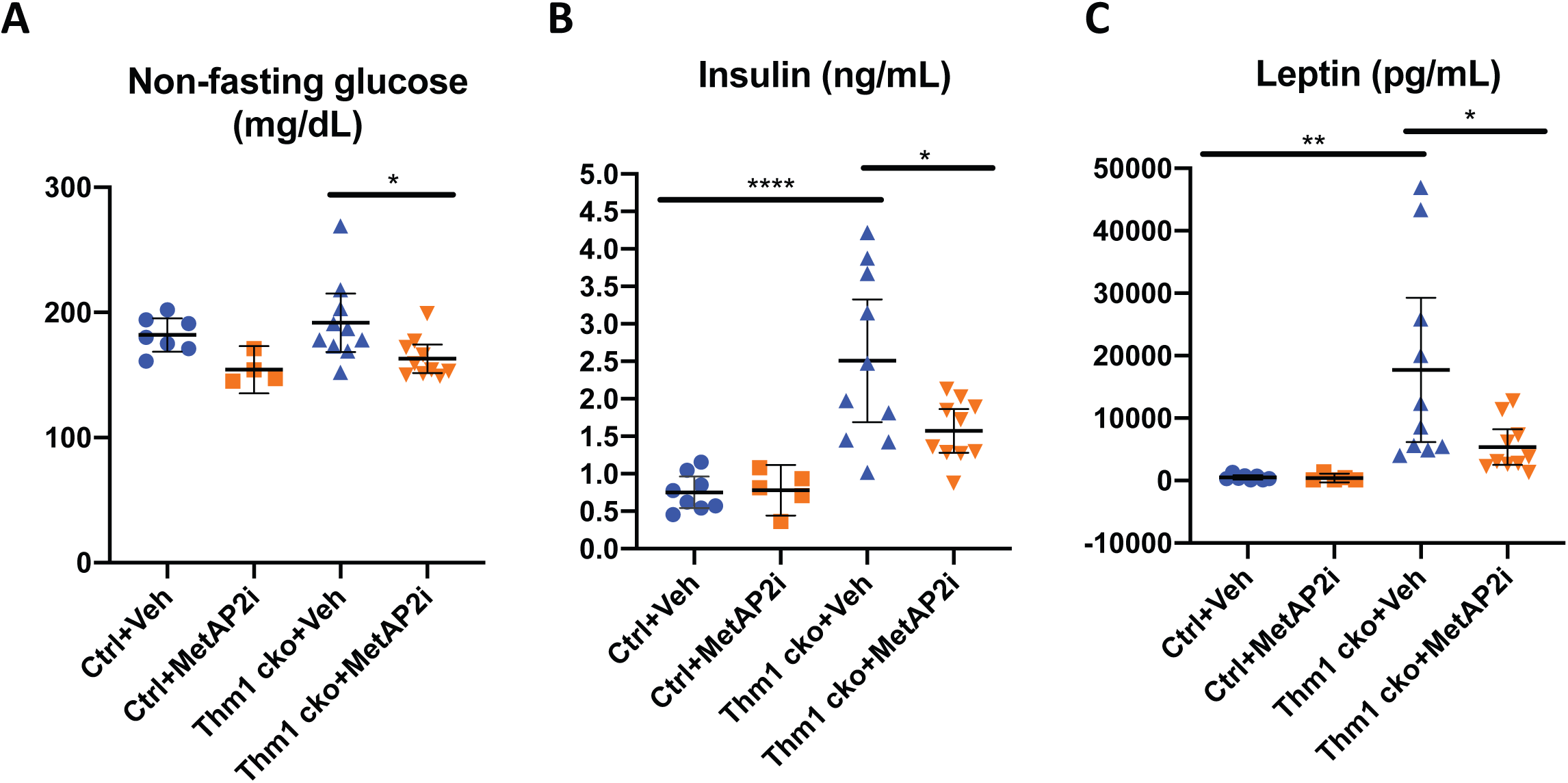
MetAP2i treatment improved metabolic parameters in *Thm1* cko mice. Non-fasting levels of **A)** blood glucose **B)** serum insulin and **C)** serum leptin. *P<0.05; **P<0.005; ****P<0.00005

### MetAP2i treatment decreases gonadal fat mass and adipocyte size in *Thm1* cko mice

We next analyzed gonadal and peri-renal fat depots, which we have shown previously to be increased in obese *Thm1* cko mice^24^. As expected, vehicle-treated *Thm1* cko mice showed increased gonadal and renal adipose tissue mass relative to control littermates (Figs 3A-D). In *Thm1* cko mice, MetAP2i treatment reduced gonadal fat mass compared to vehicle-treatment (Fig 3A). Histology of gonadal fat pads showed an increase in gonadal adipocyte cell size in vehicle-treated *Thm1* cko mice relative to control mice (Figs 3E and 3F), consistent with previous findings^24^. However, gonadal adipocyte cell size was reduced in MetAP2i-treated mutants. These data show that MetAP2i treatment partially attenuates the increased gonadal adipose tissue mass and adipocyte size caused by deletion of *Thm1*.

**Figure 3.**
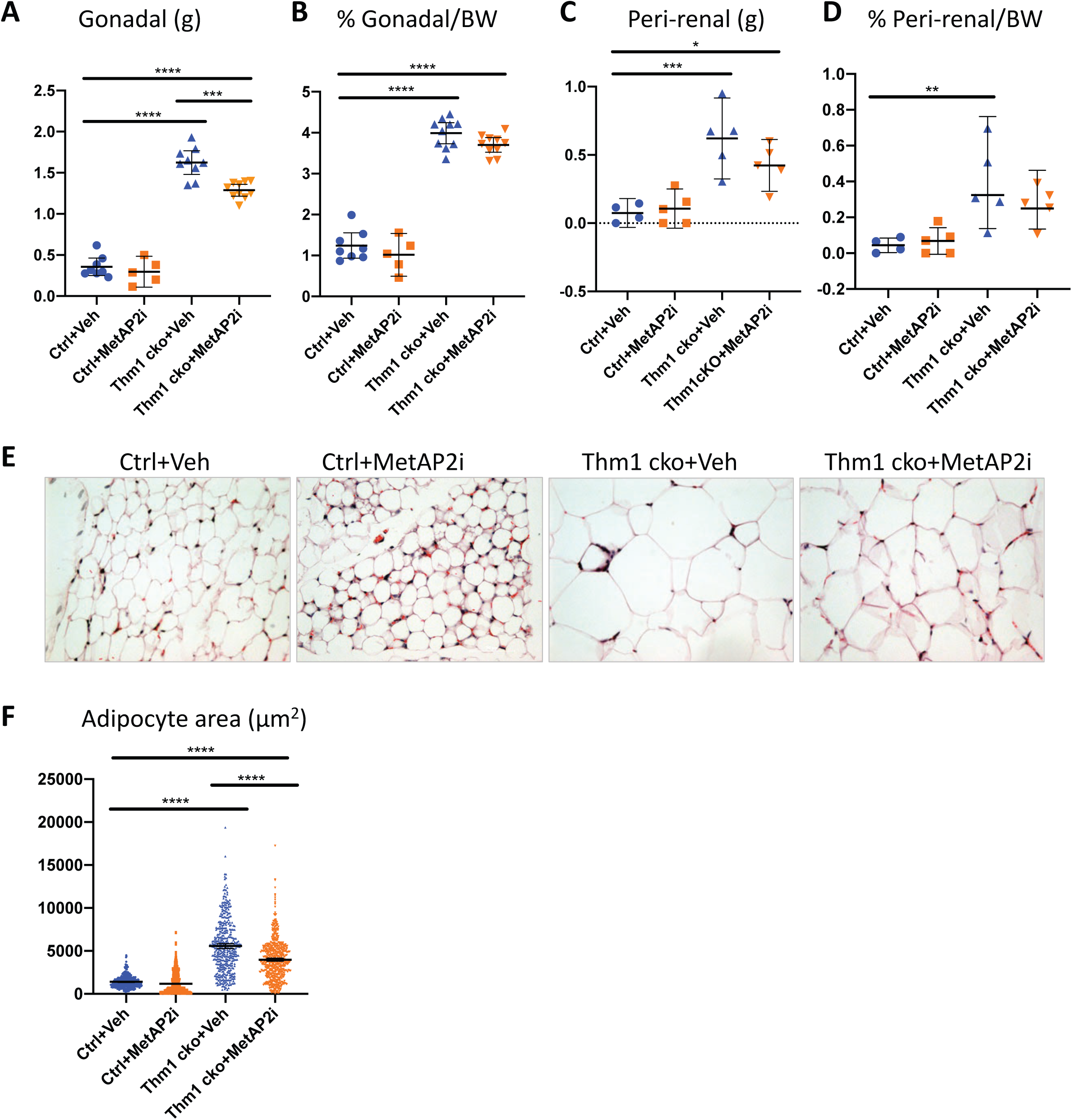
MetAP2i treatment decreased gonadal fat mass and adipocyte size in *Thm1* cko mice. **A)** gonadal adipose mass **B)** percent gonadal adipose mass/body weight **C)** peri-renal adipose mass **D)** percent peri-renal adipose mass/body weight **E)** H&E staining of gonadal adipose tissue **F)** quantification of adipocyte size. **P<0.005; ***P<0.0005; ****P<0.00005

### MetAP2i treatment improved liver morphology of *Thm1* cko mice

Obese *Thm1* cko mice can develop hepatic steatosis^24^. To determine if MetAP2i treatment affected the liver, we examined the histology of livers of vehicle- and MetAP2i-treated mice. Vehicle-treated *Thm1* cko mice had vacuoles in their livers, suggesting formation of lipid droplets (Fig 4). These vacuoles were not observed in control mice. Further, the vacuoles were reduced or absent in livers of MetAP2i-treated *Thm1* cko mice, suggesting that the improved metabolism resulting from MetAP2i treatment extends to the liver.

**Figure 4.**
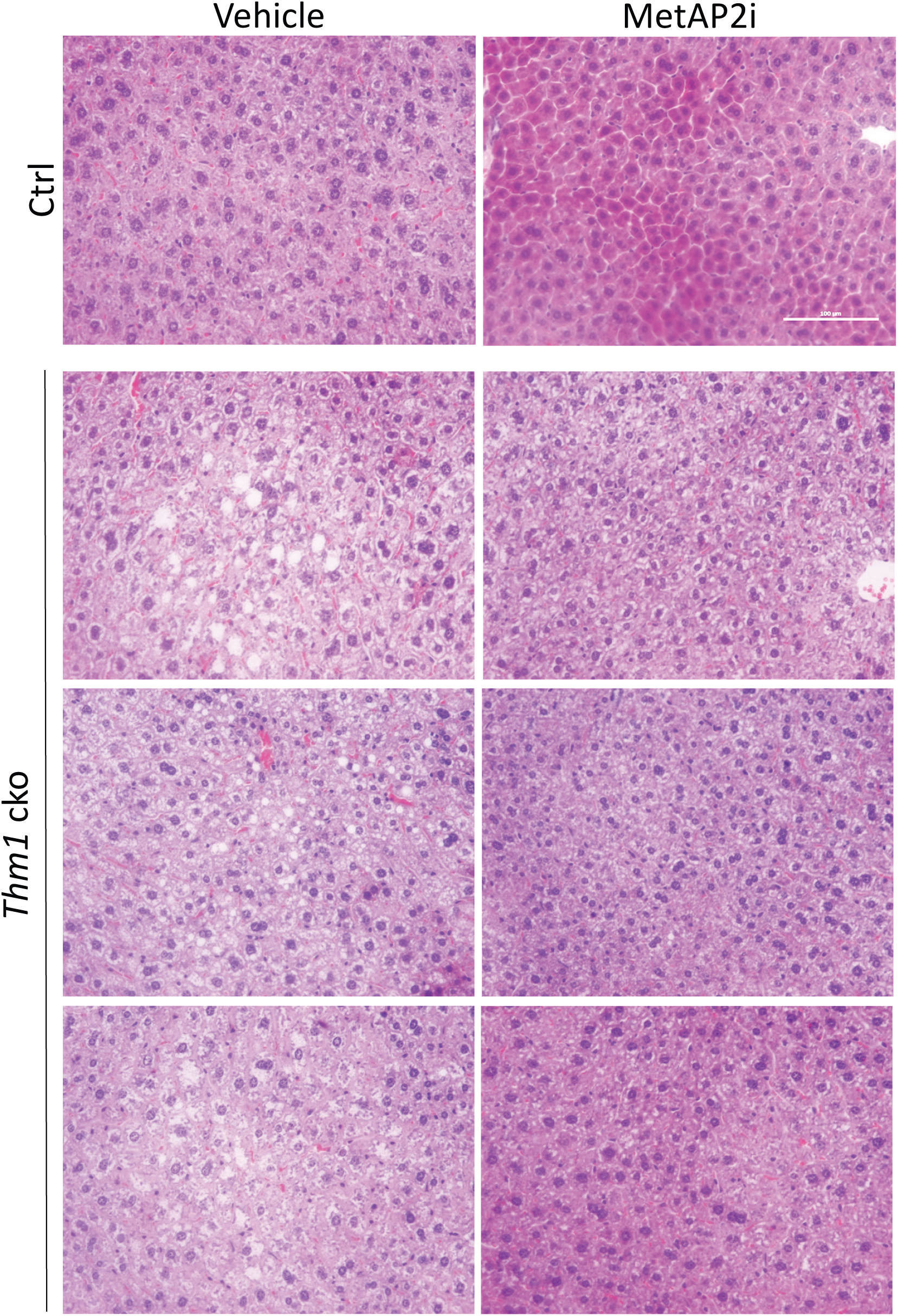
MetAP2i treatment improved liver morphology in *Thm1* cko mice. Histology of vehicle and MetAP2i-treated control (representative images) and *Thm1* cko livers. Shown are livers from 3 pairs of *Thm1* cko littermates.

### MetAP2i treatment increased renal cilia length of control and *Thm1* cko mice

Perinatal deletion of *Thm1* causes renal cystic disease by six weeks of age, but deletion of *Thm1* in adulthood does not cause renal cysts at three months following gene deletion^25^. Accordingly, kidney weights and morphology were similar between control and *Thm1* cko mice (Figs 5A and 5C). We observed a slight decrease in KW/BW ratio of vehicle-treated *Thm1* cko mice relative to vehicle-treated control mice (Fig 5B), which could be due to the increased body weight of the *Thm1* cko mice. MetAP2i treatment of control or *Thm1* cko mice did not affect kidney morphology (Fig. 5C). We next examined primary cilia by immunostaining for the ciliary membrane protein, ARL13B, together with incubating with the lectin, *Dolichos biflorus agglutinin* (DBA) to label the collecting duct. We observed previously that cilia are shortened in *Thm1*-deficient mice^24–26^. We observed that average cilia length in collecting ducts of vehicle-treated *Thm1* cko mice was shortened relative to vehicle-treated control mice, but we noted a wide range of cilia lengths in the *Thm1* cko collecting ducts and statistical significance was not achieved (Fig 1E). Unexpectedly, we observed that cilia lengths of MetAP2i-treated mice were longer than in vehicle-treated mice, and this occurred in both control and *Thm1* cko mice.

**Figure 5.**
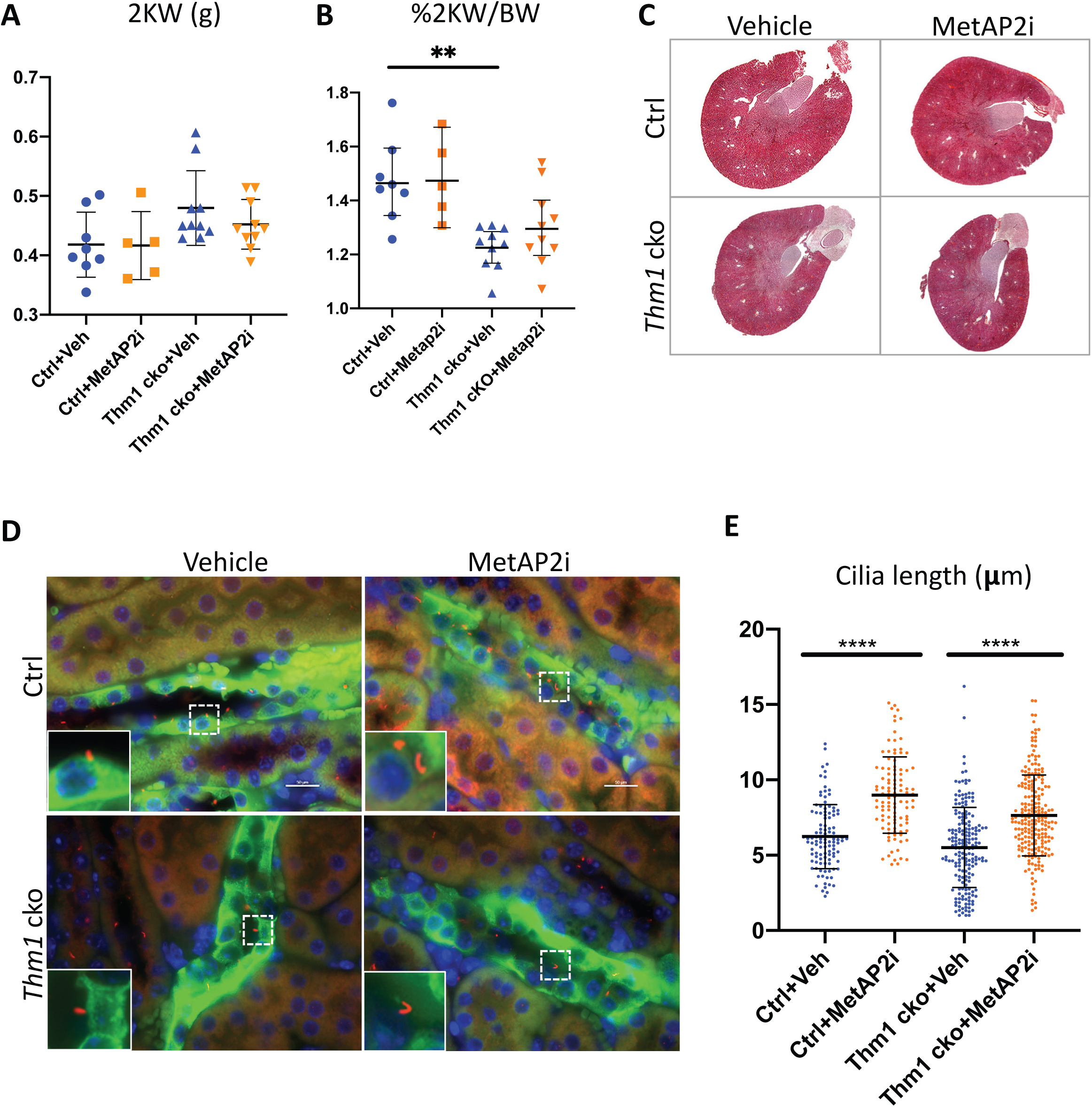
MetAP2i treatment increased renal cilia length of control and *Thm1* cko mice. **A)** 2x kidney weight **B)** Percent 2x kidney weight/body weight **C)** Histology of kidneys of vehicle and MetAP2i treated control and *Thm1* cko mice **D)** Immunostaining of primary cilia membrane protein, ARL13B (red), together with staining with lectin DBA (green), which marks the collecting duct. Insets show higher magnification of dotted boxed regions. Scale bar – 50µm. **E)** Quantification of cilia length from N=3 mice in each group. **P<0.005; ****P<0.00005

## Discussion

In this study, we provide the first evidence that MetAP2 inhibition reduces food intake and body weight and improves metabolic parameters in a ciliary model of obesity. This suggests a novel potential therapeutic strategy for ciliary disorders of obesity. This also broadens the therapeutic applicability of MetAP2 inhibition. To our knowledge, this is also the first study to show an effective pharmacological intervention in a ciliopathy rodent model of obesity.

*Thm1* deletion in adult mice results in decreased hypothalamic expression of *Pomc*, a neuropeptide which regulates appetite^24^. Reduced *Pomc* was evident before the *Thm1* cko mice gained significant body weight, suggesting reduced *Pomc* may drive the obese phenotype. Consistent with this notion, *Pomc*-null mice are obese^27^. Interestingly, in mice fed a high-fat diet, reduced hypothalamic *Pomc* expression was found to be the earliest marker predicting obesity^28^. The ability of MetAP2 inhibition to be effective in various models may indicate that MetAP2 acts on a common pathway that is misregulated in all models leading to increased appetite and obesity. Since MetAP2i reduced food intake in *Thm1* cko mice, MetAP2 may have targets in neuronal cells either downstream of or countering the effects of reduced *Pomc* expression.

Interestingly, we observed increased cilia length in renal epithelial cells of MetAP2i-treated mice. A possible mechanism by which MetAP2i increased cilia length could involve its function as an ERK inhibitor, since ERK inhibition has been shown to increase cilia length. In adipose-derived mesenchymal stem cells obtained from obese individuals, cilia are shortened^21^. However, treatment with an ERK inhibitor lengthened primary cilia and improved stemness and differentiation capacity of the cells^29^. Additionally, ERK inhibition of renal proximal tubular cells preserved cilia length during cisplatin treatment and protected the cells from cisplatin-induced apoptosis^30^. Determining in which other tissues and cells MetAP2i increased cilia length would be informative. We have shown previously that primary cilia in the *Thm1* cko hypothalamic arcuate nucleus are shortened and bulbous^24^. It seems plausible that lengthening of the mutant cilia in the arcuate nucleus may potentially attenuate the misregulated appetite. We were unable to detect primary cilia in the hypothalamus using the currently available commercial antibodies, but testing such a hypothesis in the future could reveal an important potential mechanism. Further, lengthening of primary cilia in other cell types of *Thm1* cko mice, such as adipose-derived mesenchymal stem cells, could also result in beneficial effects.

MetAP2i treatment also resulted in smaller adipocyte cell size, consistent with previous studies in rodent models of high fat diet-induced obesity^10,11^. Increased adipogenesis could result in smaller, metabolically healthier adipocytes. The direct effects of MetAP2i on adipogenesis are unclear. In one study, addition of fumagillin to an *in vitro* pre-adipocyte differentiation assay promoted adipogenesis, but had minimal effects in an *in vivo* adipogenesis assay^31^. In contrast in another study, addition of a fumagillin derivative to an *in vitro* pre-adipocyte differentiation study inhibited adipogenesis, yet fumagillin treatment of cells caused enhanced glucose uptake, suggesting metabolically healthier cells^32^. Alternatively, fumagillin has anti-angiogenic effects and a prevailing hypothesis is that changes in angiogenesis induced by fumagillin reduce adipose tissue mass. Yet, a study showed that angiogenesis changes did not drive reduction of adipose tissue mass^10^. Thus, further studies are required to determine the mechanisms by which adipocyte size and adipose tissue mass are reduced by MetAP2i.

Given safety issues, Zafgen has recently discontinued the development of MetAP2 inhibitors^33^. Our study indicates that there are plausible targets of MetAP2 inhibition that effectively reduce hyperphagia and body weight and substantially improve metabolic parameters in a ciliopathy model. This substantiates the need for greater understanding of the biology of MetAP2^34^. Currently, agonists of the melanocortin 4 receptor, which is activated by the processed protein products of *Pomc*, are being tested in clinical trial to target obesity in patients with BBS and Alström Syndrome^35^. Aside from this, a potential therapy for ciliopathy-induced obesity has not been demonstrated. The mechanism by which MetAP2 inhibition exerts its anti-obesity effects remain elusive. Intriguingly, our data suggest MetAP2 inhibition modifies cilia length irrespective of genotype, and therefore, may alter cilia length in other models of obesity. Whether changes in ciliary dynamics underlie the beneficial effects of MetAP2 inhibition may warrant investigation, and could reveal the therapeutic targets.

## Methods

### Generation of *Thm1* cko mice

*Thm1* cko mice were generated as described^25^. Briefly, *Thm1* cko mice were generated by using a *Thm1* null allele (called *aln*) and a floxed allele, which has LoxP sites flanking exon 4, together with a tamoxifen-inducible *ROSA26-CreERT* recombinase (Jackson Laboratories, Stock 004847), which is expressed globally. Cre recombinase expression was induced at 5 weeks of age by i.p. injection of 10 mg tamoxifen/40g mouse weight. Tamoxifen (Sigma T5648) was suspended in corn oil (Sigma C8267) at 30 mg/ml. Only male mice were used.

### Administration of MetAP2i

Mice were fed *ab libitum* throughout the duration of the study. Body weight was measured weekly from 0-10 weeks post-tamoxifen injection to ascertain the obese phenotype in *Thm1* cko mice. Beginning at 10 weeks post-tamoxifen injection, mice were housed individually, and food intake and body weight were measured daily until the end of the experiment. Subcutaneous injections of MetAP2i (ZGN-1258 - 0.3 mg/kg/day) or saline vehicle were administered daily from 11-13 weeks post-tamoxifen injection. All animal studies were approved by KUMC IACUC.

### Blood glucose and serum insulin and leptin measurements

Blood glucose was measured from tail blood using Bayer Blood Glucose Contour Strips together with the Bayer Contour Blood Glucose Meter system. For serum insulin and leptin measurements, trunk blood was collected in a Microvette CB300z blood collection tube (Kent Scientific), and serum was isolated by centrifuging blood collection tubes for 6 minutes at 800xg at 4°C using a tabletop centrifuge (PrismR, C2500-R). Insulin and leptin levels were measured using Mouse Ultrasensitive Insulin ELISA and Mouse/Rat Leptin ELISA kits (ALPCO) according to manufacturer’s instructions.

### Histology

Tissues were isolated and submerged in 10% formalin for 3-7 days. Fixed tissues were dehydrated through an ethanol series, paraffin-embedded and sectioned at 7µm thicknesses. Sections were rehydrated and stained with hematoxylin and eosin using a standard protocol. Staining was viewed and imaged using a Nikon 80i microscope equipped with a Nikon DS-Fi1 camera.

### Immunofluorescence

Immunofluorescence was performed as described (Silva et al., 2019). Sections were rehydrated. Heat antigen retrieval was performed in tri-sodium citrate solution, pH 6.0, using a steamer. Sections were rinsed in distilled water (10X), then blocked in 2% BSA in PBS for 1 hour. Arl13B antibody (Proteintech) was diluted in blocking buffer (2% BSA in PBS) and tissue sections were incubated with primary antibody at 4°C overnight. Sections were washed 3X in PBS, then incubated with *Dolichos biflorus agglutinin* (DBA; Vector Laboratories) for 1 hour at room temperature. Sections were washed in PBS 3X, then incubated with Alexa Fluor 594, goat anti-rabbit secondary antibody (ThermoFisher) for 1 hour at room temperature. Following 3 washes in PBS, tissue sections were then mounted with Fluoromount G containing DAPI mounting media (Electron Microscopy Services). Immuno-labeled tissues were viewed and imaged using a Leica TCS SPE confocal microscope configured on a DM550 Q upright microscope.

### Statistics

Graphpad Prism 8 was used to perform statistical analyses, which included two-tailed unpaired t-tests, and ANOVA followed by Tukey’s test.

## Acknowledgements

We thank members of the KUMC Department of Anatomy and Cell Biology and of the Kidney Institute for helpful discussions. We also thank the KUMC KIDDRC Core Services - Jing Huang of the Histology Core and Michelle Winter of the Animal Behavioral Core. Work performed by the Cores is supported by the KUMC Smith Intellectual and Developmental Disabilities Research Center (NIH U54 HD 090216).

## References

1. Roth J, Qiang X, Marban SL, Redelt H, Lowell BC. The obesity pandemic: where have we been and where are we going? Obes Res. 2004;12 Suppl 2:88S–101S.

2. Lowther WT, Matthews BW. Structure and function of the methionine aminopeptidases. Biochim Biophys Acta. 2000;1477(1-2):157–167.

3. Liu S, Widom J, Kemp CW, Crews CM, Clardy J. Structure of human methionine aminopeptidase-2 complexed with fumagillin. Science. 1998;282(5392):1324–1327.

4. Turk BE, Griffith EC, Wolf S, Biemann K, Chang YH, Liu JO. Selective inhibition of amino-terminal methionine processing by TNP-470 and ovalicin in endothelial cells. Chem Biol. 1999;6(11):823–833.

5. Yeh JR, Mohan R, Crews CM. The antiangiogenic agent TNP-470 requires p53 and p21CIP/WAF for endothelial cell growth arrest. Proc Natl Acad Sci U S A. 2000;97(23):12782–12787.

6. Datta B, Datta R, Majumdar A, Ghosh A. The stability of eukaryotic initiation factor 2-associated glycoprotein, p67, increases during skeletal muscle differentiation and that inhibits the phosphorylation of extracellular signal-regulated kinases 1 and 2. Exp Cell Res. 2005;303(1):174–182.

7. Shimizu H, Yamagishi S, Chiba H, Ghazizadeh M. Methionine Aminopeptidase 2 as a Potential Therapeutic Target for Human Non-Small-Cell Lung Cancers. Adv Clin Exp Med. 2016;25(1):117–128.

8. Dipaolo JA, Tarbell DS, Moore GE. Studies on the carcinolytic activity of fumagillin and some of its derivatives. Antibiot Annu. 1958;6:541–546.

9. Garvey MJ, Ambrose PG, Ulmer JL. Topical fumagillin in the treatment of microsporidial keratoconjunctivitis in AIDS. Ann Pharmacother. 1995;29(9):872–874.

10. Lijnen HR, Frederix L, Van Hoef B. Fumagillin reduces adipose tissue formation in murine models of nutritionally induced obesity. Obesity. 2010;18(12):2241–2246.

11. An J, Wang L, Patnode ML, et al. Physiological mechanisms of sustained fumagillin-induced weight loss. JCI Insight. 2018;3(5).

12. Brakenhielm E, Cao R, Gao B, et al. Angiogenesis inhibitor, TNP-470, prevents diet-induced and genetic obesity in mice. Circ Res. 2004;94(12):1579–1588.

13. Hughes TE, Kim DD, Marjason J, Proietto J, Whitehead JP, Vath JE. Ascending dose-controlled trial of beloranib, a novel obesity treatment for safety, tolerability, and weight loss in obese women. Obesity. 2013;21(9):1782–1788.

14. McCandless SE, Yanovski JA, Miller J, et al. Effects of MetAP2 inhibition on hyperphagia and body weight in Prader-Willi syndrome: A randomized, double-blind, placebo-controlled trial. Diabetes Obes Metab. 2017;19(12):1751–1761.

15. Baker K, Beales PL. Making sense of cilia in disease: the human ciliopathies. Am J Med Genet C Semin Med Genet. 2009;151C(4):281–295.

16. Mukhopadhyay S, Wen X, Chih B, et al. TULP3 bridges the IFT-A complex and membrane phosphoinositides to promote trafficking of G protein-coupled receptors into primary cilia. Genes Dev. 2010;24(19):2180–2193.

17. Fu W, Wang L, Kim S, Li J, Dynlacht BD. Role for the IFT-A Complex in Selective Transport to the Primary Cilium. Cell reports. 2016;17(6):1505–1517.

18. Guo DF, Rahmouni K. Molecular basis of the obesity associated with Bardet-Biedl syndrome. Trends in endocrinology and metabolism: TEM. 2011;22(7):286–293.

19. Girard D, Petrovsky N. Alstrom syndrome: insights into the pathogenesis of metabolic disorders. Nature reviews Endocrinology. 2011;7(2):77–88.

20. Benzinou M, Walley A, Lobbens S, et al. Bardet-Biedl syndrome gene variants are associated with both childhood and adult common obesity in French Caucasians. Diabetes. 2006;55(10):2876–2882.

21. Ritter A, Friemel A, Kreis NN, et al. Primary Cilia Are Dysfunctional in Obese Adipose-Derived Mesenchymal Stem Cells. Stem Cell Reports. 2018;10(2):583–599.

22. Ritter A, Louwen F, Yuan J. Deficient primary cilia in obese adipose-derived mesenchymal stem cells: obesity, a secondary ciliopathy. Obesity reviews: an official journal of the International Association for the Study of Obesity. 2018;19(10):1317–1328.

23. Davis EE, Zhang Q, Liu Q, et al. TTC21B contributes both causal and modifying alleles across the ciliopathy spectrum. Nat Genet. 2011;43(3):189–196.

24. Jacobs DT, Silva LM, Allard BA, et al. Dysfunction of intraflagellar transport-A causes hyperphagia-induced obesity and metabolic syndrome. Dis Model Mech. 2016;9(7):789–798.

25. Tran PV, Talbott GC, Turbe-Doan A, et al. Downregulating hedgehog signaling reduces renal cystogenic potential of mouse models. J Am Soc Nephrol. 2014;25(10):2201–2212.

26. Tran PV, Haycraft CJ, Besschetnova TY, et al. THM1 negatively modulates mouse sonic hedgehog signal transduction and affects retrograde intraflagellar transport in cilia. Nat Genet. 2008;40(4):403–410.

27. Yaswen L, Diehl N, Brennan MB, Hochgeschwender U. Obesity in the mouse model of pro-opiomelanocortin deficiency responds to peripheral melanocortin. Nat Med. 1999;5(9):1066–1070.

28. Souza GF, Solon C, Nascimento LF, et al. Defective regulation of POMC precedes hypothalamic inflammation in diet-induced obesity. Scientific reports. 2016;6:29290.

29. Ritter A, Kreis NN, Roth S, et al. Restoration of primary cilia in obese adipose-derived mesenchymal stem cells by inhibiting Aurora A or extracellular signal-regulated kinase. Stem Cell Res Ther. 2019;10(1):255.

30. Wang S, Wei Q, Dong G, Dong Z. ERK-mediated suppression of cilia in cisplatin-induced tubular cell apoptosis and acute kidney injury. Biochim Biophys Acta. 2013;1832(10):1582–1590.

31. Scroyen I, Christiaens V, Lijnen HR. Effect of fumagillin on adipocyte differentiation and adipogenesis. Biochim Biophys Acta. 2010;1800(4):425–429.

32. Siddik MAB, Das BC, Weiss L, Dhurandhar NV, Hegde V. A MetAP2 inhibitor blocks adipogenesis, yet improves glucose uptake in cells. Adipocyte. 2019;8(1):240–253.

33. ZAFGEN TO EXPLORE STRATEGIC ALTERNATIVES: Preliminary results from in vivo animal study not expected to warrant near-term resolution of clinical hold for ZGN-1061 [Press release] [press release]. https://zafgen.gcs-web.com/news-releases/news-release-details/zafgen-explore-strategic-alternatives, September 5, 2019 2019.

34. Chang YH. Common therapeutic target for both cancer and obesity. World J Biol Chem. 2017;8(2):102–107.

35. Eneli I, Xu J, Webster M, et al. Tracing the effect of the melanocortin-4 receptor pathway in obesity: study design and methodology of the TEMPO registry. Appl Clin Genet. 2019;12:87–93.

