## Supplementary figures and images for "MetAP2 inhibition reduces food intake and body weight in a ciliopathy mouse model of obesity"

### Supplemental Figure 1

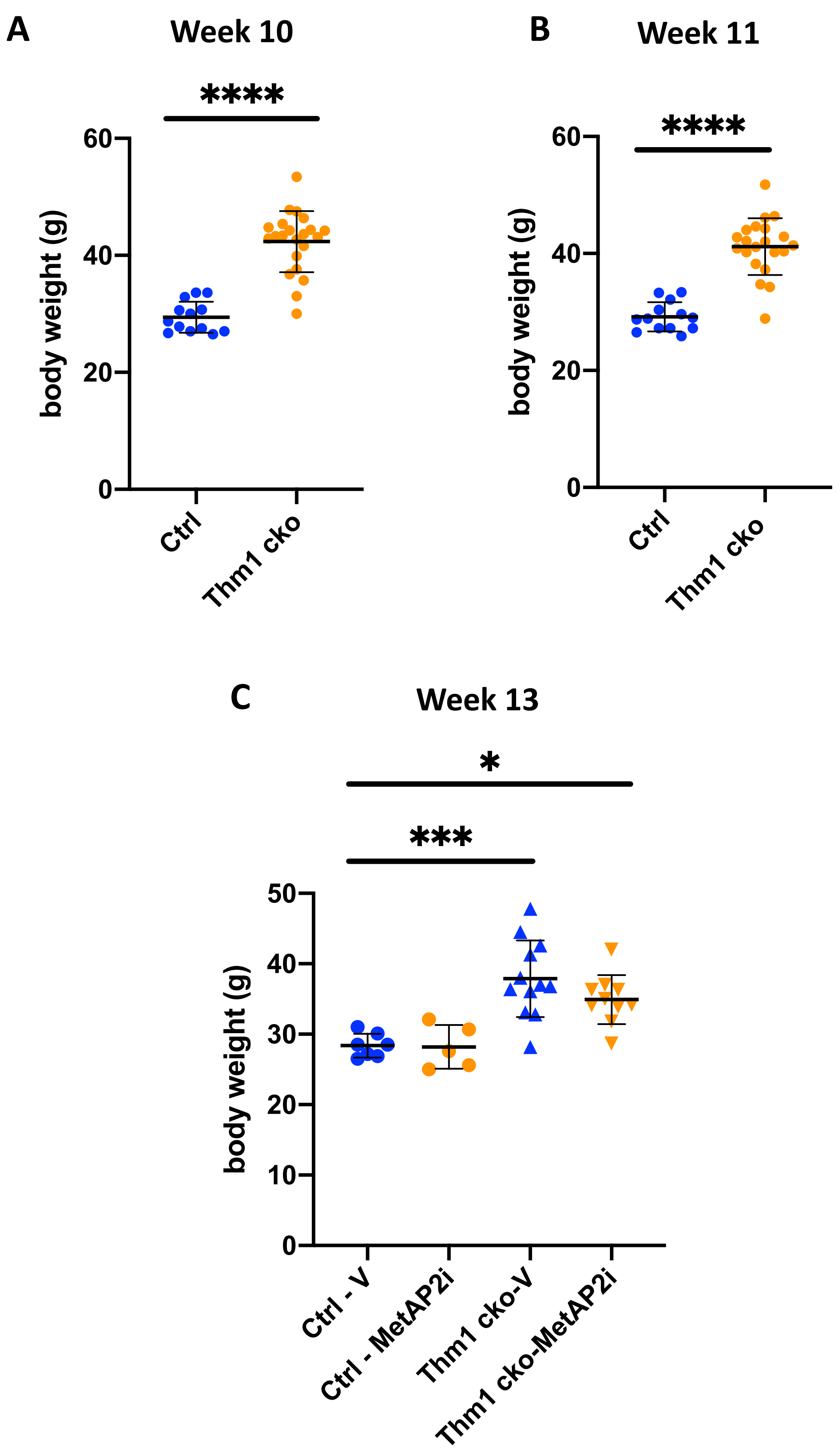


**Figure S1. Body weights of control and *Thm1* cko mice. A)** week 10 **B)** week 11 **C)** week 13.
*P<0.05; ***P<0.0005; ****P<0.00005
